# Designing a Model to Detect Beta Burst in EEG Using Nonlinear Dynamic Features Based on Machine Learning

**DOI:** 10.1101/2025.10.13.682006

**Authors:** Armin Hakkak Moghadam Torbati, Narges Davoudi, Giuseppe Longo

## Abstract

**Background:** Beta bursts are brief, transient increases in beta-band (13–30 Hz) EEG activity that play a key role in motor control, particularly in processes like movement initiation and inhibition. While most existing methods detect these bursts using simple amplitude thresholds, they often ignore variability in burst duration and task context. More advanced techniques exist but are computationally demanding, often opaque, and dependent on large datasets and strong modeling assumptions.

**Objectives:** This study aims to develop an automated, machine learning–based approach to classify beta bursts using nonlinear dynamic features, considering both burst duration and task condition.

**Method:** EEG data were collected from 26 healthy, right-handed participants during three motor tasks: (1) a right-hand isometric pinch grip at 10% maximum voluntary contraction (MVC), (2) rhythmic right-hand finger opening–closing in response to auditory cues (3–4 seconds), and (3) the same right-hand opening–closing task and concurrent with a steady left-hand isometric pinch grip at 10% MVC. Beta bursts were extracted from the left motor cortex, categorized by duration (short, medium, long), and time-locked to task events. From each burst, four nonlinear features, Fractal Dimension (FD), Wavelet Entropy (WE), Sample Entropy (SE), and Nonlinear Energy Operator (NEO) were calculated to train machine learning (ML) models.

**Results:** Statistical tests and feature selection revealed that FD, SE, and WE varied significantly with burst duration and task type, while NEO was more limited in sensitivity. ML models trained on these features achieved up to 91.1% validation and 85.7% test accuracy, especially when distinguishing bursts of different durations within the same task.

**Conclusions:** These findings suggest that beta bursts reflect structured, task-specific neural dynamics rather than random fluctuations. By using nonlinear features and ML, we introduce a scalable, interpretable framework for burst classification that outperforms traditional methods. This approach advances the understanding of transient beta activity and holds promise for real-time neural decoding in neuroscience and neurotechnology.

## 1. Introduction

Beta oscillations (13–30 Hz) play a well-established role in motor control and have traditionally been viewed as continuous signals linked to movement preparation and execution [1, 2]. While this perspective has yielded valuable insights, it largely overlooks the transient nature of beta activity. Recently, research has shifted focus toward brief, intermittent bursts of beta activity, offering a more nuanced understanding of neural dynamics [3, 4]. These beta bursts are now increasingly recognized as functionally significant, reflecting critical aspects of brain processing rather than mere background noise [5].

In the motor domain, the timing and duration of beta bursts have been associated with key processes such as movement initiation, inhibition, cancellation, and error monitoring [6, 7]. For example, movement initiation is typically preceded by a reduction in beta bursting across bilateral sensorimotor areas, whereas rapid movement cancellation elicits bursts over fronto-central motor control regions, followed by sensorimotor reactivation [6]. In clinical populations such as Parkinson’s disease (PD), prolonged beta bursts are linked to motor rigidity and slowness, whereas shorter, well-timed bursts facilitate smoother movements [8]. These observations suggest that beta bursts modulate motor behavior rather than merely reflect ongoing motor output.

Despite the growing recognition of beta bursts, many analytical methods continue to treat them as uniform events. Traditional threshold-based detection [9], is simple and practical but fails to capture variability in burst shape, duration, or task-dependent modulation. More advanced techniques—such as adaptive waveform modeling and dimensionality reduction (e.g., PCA on waveform motifs)—offer richer insights but are computationally demanding, often opaque, and dependent on large datasets and strong modeling assumptions [10, 11]. Therefore, there remains a need for a computationally efficient method to capture beta bursts reliably and rapidly.

The utilization of nonlinear dynamic features of beta bursts may serve as a feasible methodological approach. This is because beta oscillations typically occur as transient bursts consisting of a few cycles separated by silent periods, an activity pattern that reflects the complex and inherently nonlinear dynamics of neural systems [12]. The relevance of such dynamic characteristics in brain oscillations has been demonstrated across numerous studies, ranging applications from the diagnosis of neurodegenerative disorders to the assessment of cognitive and neural states [13, 14]. Moreover, the rapid and automated detection and classification of beta bursts hold considerable potential for many applications such as brain– computer interface (BCI) [15]. A particularly suitable strategy involves the integration of machine learning (ML) methods, which offer both speed and robustness, enabling real-time burst detection. Accordingly, we hypothesize that combining nonlinear dynamic feature analysis with ML-based classification may provide an effective framework for detecting and categorizing beta bursts both across different motor tasks and burst duration. In fact, ML algorithms can capture complex patterns within nonlinear features and generalize effectively across datasets, making them powerful tools for automated beta-burst analysis.

This study investigates whether nonlinear dynamic features can effectively characterize beta bursts with respect to both duration and motor task condition. Specifically, we extract features such as fractal dimension (FD), wavelet entropy (WE), nonlinear energy operator (NEO), and sample entropy (SE) from EEG-detected beta bursts, and analyze their variation across short, medium, and long bursts and across distinct motor tasks. We further develop ML models to classify beta bursts based on these nonlinear features, aiming to differentiate burst types by both duration and task-related dynamics. This approach aims to provide a fast, automatic, interpretable, and functionally meaningful framework for beta-burst analysis.

To address these goals, we propose two research questions:

1. Do nonlinear dynamic features (FD, WE, NEO, SE) differ significantly across beta bursts of different durations (short, medium, long) and across motor task conditions? Hypothesis 1: Nonlinear features will show statistically significant variation with burst duration and task type, reflecting differences in underlying neural dynamics.
2. Can an ML model accurately classify beta burst types (by duration and task condition) based on extracted nonlinear features?

Hypothesis 2: Nonlinear features contain sufficient discriminative information to enable accurate ML-based classification of burst types.

## 2. Method and materials

### 2.1. Participants

Twenty-six healthy, right-handed adult volunteers (14 males, 12 females; mean age: 25 ± 5 years; range: 18–30) participated in the study. All participants had no history of neurological or psychiatric disorders and provided written informed consent prior to participation. The study was approved by the Ethics Committee of Erasme Hospital, Université Libre de Bruxelles (Belgium), and participants received monetary compensation for travel and time.

### 2.2. Experimental Protocol

The experiment was conducted in a quiet, controlled laboratory setting. Participants sat comfortably with both forearms supported in a symmetrical position. Each participant performed the following three motor tasks:

1. **Right-hand isometric task**: Participants maintained a steady isometric pinch grip at 10% of their maximum voluntary contraction (MVC) using the right hand.
2. **Right-hand dynamic task**: Participants opened and closed their right-hand fingers in response to beep sounds played every 3–4 seconds with random jitter.
3. **Bimanual dual task**: Participants performed the same right-hand opening–closing task as in Task 2, while simultaneously maintaining a steady isometric pinch grip at 10% of their MVC with the left hand (fig 1).

**Fig. 1.**
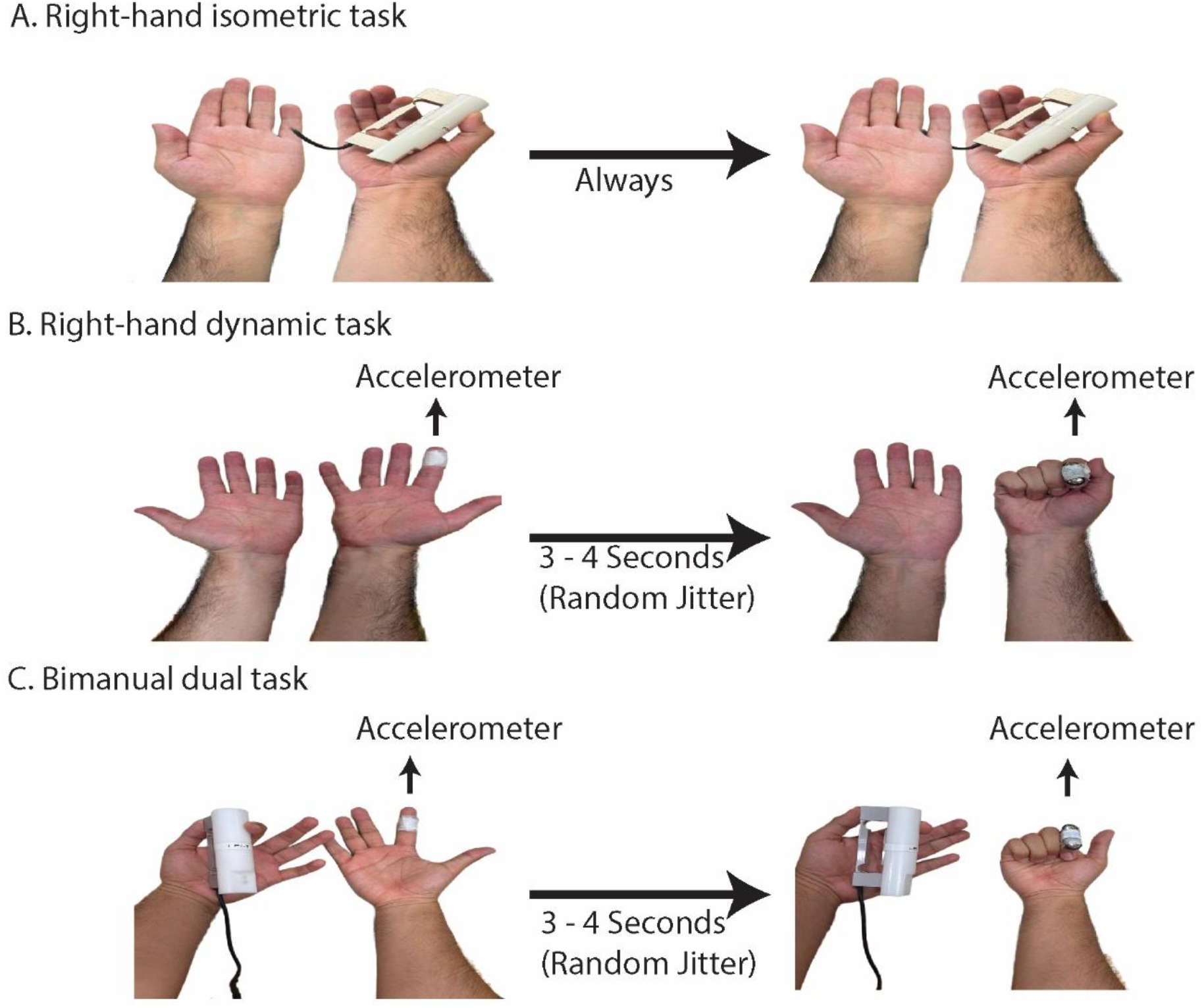
Task design and experimental conditions. **A)** Right-hand isometric task: Participants maintained a steady 10% MVC pinch grip using a custom-built force transducer throughout the trial. **B)** Right-hand dynamic task: Participants repeatedly opened and closed their right hand in response to auditory cues played every 3–4 seconds (3–4 s intervals with random jitter). An accelerometer on the index finger detected movement onset. **C)** Bimanual dual task: In the bimanual dual-task condition, the same right-hand movements were performed concurrently with a steady left-hand isometric pinch grip maintained at 10 % of the participant’s MVC.

Each task condition lasted 5 minutes and was repeated twice, with at least 2 minutes of rest between blocks. MVC was assessed bilaterally using a hand dynamometer (range: 0–500 N). During tasks requiring force control, participants received real-time visual feedback (Fig 2) through a custom MATLAB interface to maintain the target force level. The experimental setup is depicted in Fig 1.

**Fig. 2.**
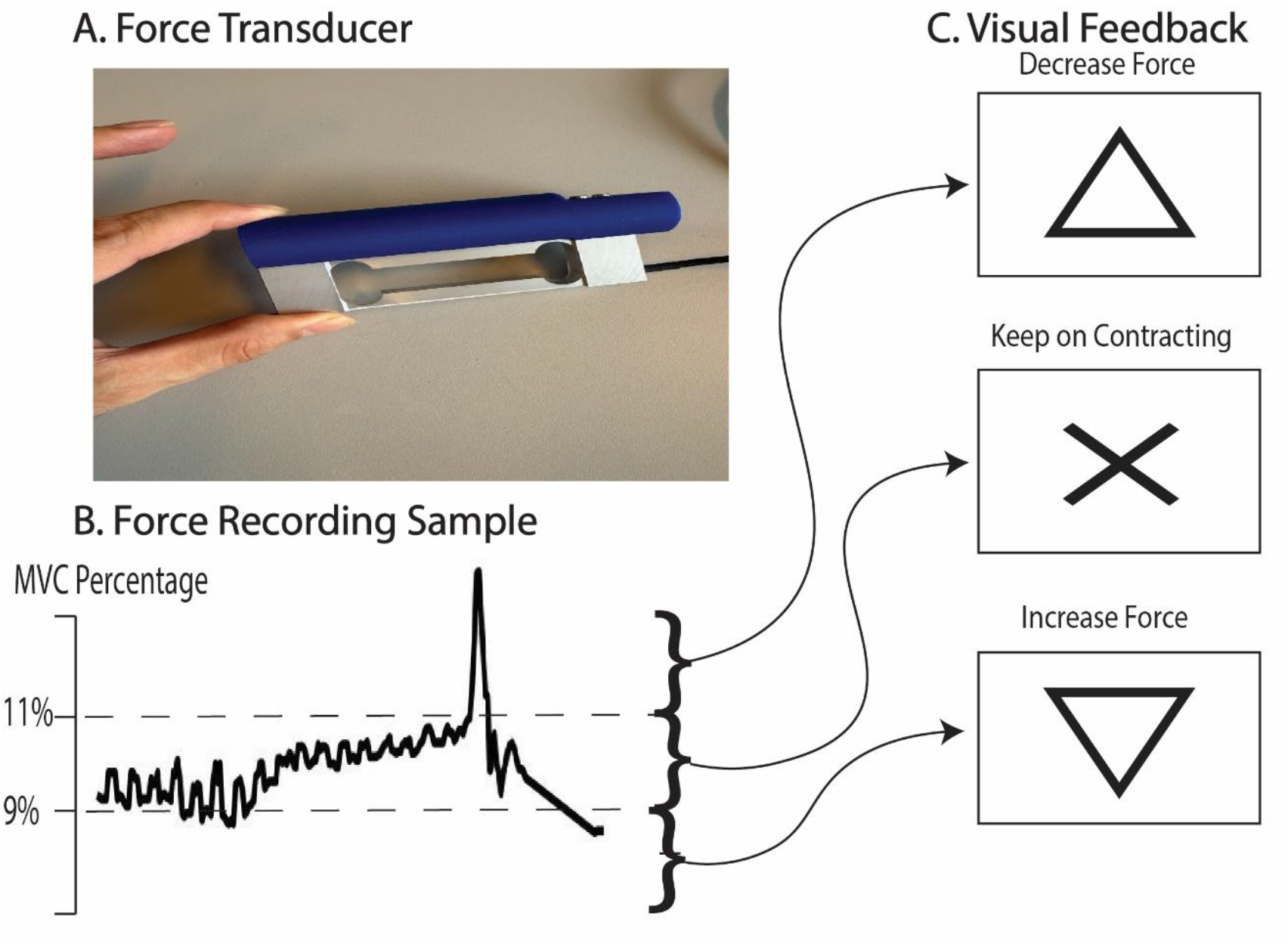
Visual feedback interface for controlling isometric force. **A)** Custom-built force transducer used for right-hand pinch grip, positioned between the thumb and index finger. The device was designed to measure steady isometric contractions at a target force level. **B)** Sample trace of recorded force output from a participant. Dashed horizontal lines indicate the acceptable force range (9–11% of maximum voluntary contraction, MVC). Deviations outside this range triggered visual feedback. **C)** Real-time visual cues provided to guide force regulation. A central cross was shown when the force remained within bounds. An upward- or downward-pointing triangle was displayed when force exceeded or dropped below the threshold, prompting the participant to decrease or increase their grip force, respectively.

### 2.3. Measurement

EEG signals were recorded using a 64-channel EEG system (EEGO Mylab, ANT Neuro, Netherlands) following the extended 10–20 system. Electrode impedance was maintained below 20 kΩ, and signals were sampled at 1 kHz with CPz as the reference. A 3-axis accelerometer (COMETA Pico, Milan, Italy) was attached to the right-hand index finger to record movement onset during dynamic tasks (task 2 and 3). An additional sensor under the eye was used to monitor artifacts. In Tasks 1 and 3, isometric force was exerted on a custom transducer (range: 0–6 N) incorporating a model 1004 load cell (Vishay Precision Group, USA), and recorded simultaneously via the COMETA Pico system (COMETA, Milan, Italy) to ensure synchronization with EEG. EMG signal was also recorded from the First Dorsal Interosseous (rFDI) muscle with this system.

### 2.4. Preprocessing

EEG data preprocessing was conducted offline using custom MATLAB scripts (MathWorks, Natick, MA) and the FieldTrip toolbox [16]. Bad channels were identified according to the criteria described by Bigdely-Shamlo et al [17] and subsequently interpolated using topographic methods. The EEG signals were re-referenced to a common average and band-pass filtered between 0.3 and 45 Hz. To further eliminate artifacts, independent component analysis (ICA) was applied following the approach described by Vigário et al. [18]. Prior to ICA, the data dimensionality was reduced to 25 principal components using principal component analysis (PCA) to improve numerical stability. ICA was then computed using the FastICA algorithm with a hyperbolic tangent nonlinearity [18, 19]. From this decomposition, 20 independent components were retained for inspection. Components associated with ocular and cardiac artifacts, such as eye blinks, eye movements, and heartbeat activity, were identified through visual inspection of their scalp topographies, time courses, and power spectra and subsequently removed. On average, 3 components per participant (SD = 1.2) were excluded. The remaining components were back-projected to sensor space for subsequent beta-burst detection and feature extraction.

EEG, force, EMG, and accelerometer signals were temporally synchronized using a digital trigger generated by the experimental control PC and recorded simultaneously by all acquisition systems. This trigger was used solely to ensure precise temporal alignment across modalities and was not used to define analysis epochs. For experiments involving movement (experiments 2 and 3), EEG epochs were time-locked to movement onset detected from accelerometer signals, whereas for experiment 1, the continuous EEG signal was segmented into fixed-length windows to match epoch duration across tasks. Figure 3 illustrates a representative example of the synchronized EMG signal with digital trigger markers, highlighting the common temporal reference used across modalities. All signals were filtered using a low-pass cosine filter with a cutoff frequency of 50 Hz, implemented in the frequency domain, to reduce high-frequency noise and electrical interference. In experiments 1 and 3, periods during which the force deviated from the target 10% MVC range, based on visual feedback, were excluded from the force signal as well as the corresponding EEG segments.

**Fig. 3.**
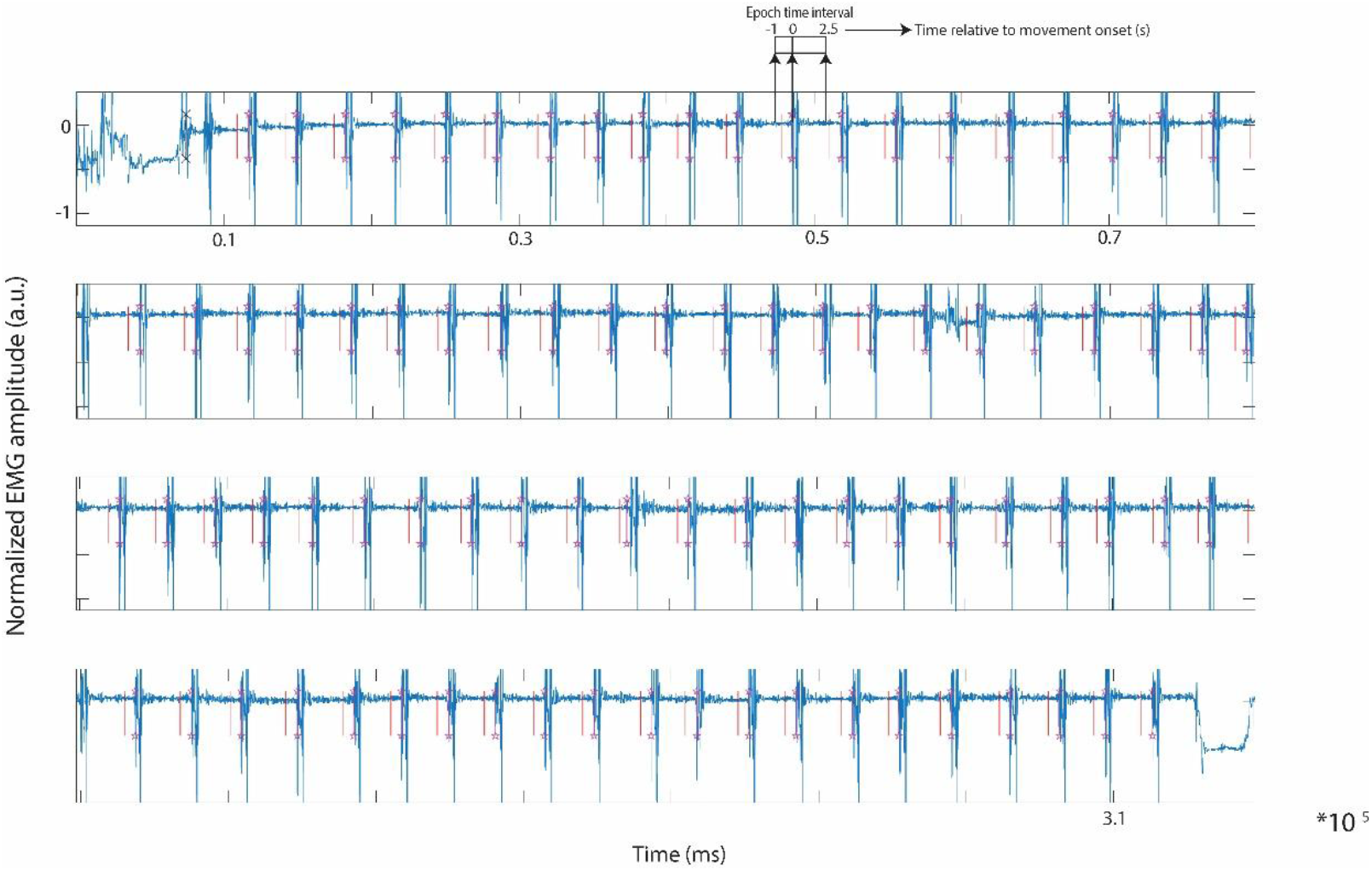
Representative example of synchronized EMG signals, event markers, and EEG observation windows. Representative EMG recordings from a single subject are shown across consecutive epochs. Vertical red lines indicate digital trigger markers used for temporal synchronization between EEG, EMG, and accelerometer recordings. Purple star markers denote movement-related events detected from the accelerometer signal and used for time-locking epochs in movement-related experiments. Arrows indicate the EEG observation window (−1 to +2.5 s) defined relative to movement onset and used for beta-burst analysis, as described in Section 2.5. The figure illustrates the common temporal reference across modalities and the selection of analysis windows.

### 2.5 Beta Burst Detection and Nonlinear Dynamic Features Extraction

To identify and characterize beta bursts from EEG signals, we processed preprocessed data recorded from electrodes over the left motor cortex (channels: ‘C1’ ‘C3’ ‘C5’ ‘CP1’ ‘CP3’ ‘CP5’ ‘FC1’ ‘FC3’ ‘FC5’). The averaged signal was bandpass filtered in the beta range (13–30 Hz), and the amplitude envelope was extracted using the Hilbert transform. To normalize for slow fluctuations, the envelope was divided by a smoothed version filtered below 0.1 Hz. For experiments 2 and 3, trials were time-locked to movement onset, detected via accelerometer signals, and segmented from −1000 ms to +2500 ms. But in the first experiment, in the absence of movement events, the continuous signal was divided into 3500-ms time windows to match the epoch length across all tasks. We excluded epochs with unusually large signal amplitudes, defined as those more than five interquartile ranges (IQRs) above the median. Beta bursts were defined as periods when the amplitude was greater than the 75th percentile of the overall amplitude distribution.

Only bursts lasting at least 50 samples (≈50 ms) were considered valid. Bursts were categorized into short (<100 ms), medium (100–149 ms), and long (≥150 ms) durations. This segmentation is based on prior studies showing that beta bursts exhibit substantial variability in duration, and burst duration is linked to functional relevance rather than reflecting a uniform oscillatory process. In particular, very short beta bursts (typically <100 ms) are often interpreted as brief, transient events, whereas longer bursts (≥150 ms) have been associated with more sustained cortical engagement and greater relevance for motor control and behavioural state [5, 20, 21]. Rather than assuming discrete functional roles for each duration category, the present study adopted short, medium, and long bins to capture these broad temporal regimes while allowing for intermediate-duration events. These boundaries were further informed by the empirical distribution of burst durations in the present dataset, which exhibited natural separation between short-lived and longer-lasting events. Importantly, the selected bins represent broad duration ranges rather than precise cut-offs, and modest shifts in boundary values are therefore unlikely to qualitatively alter the reported statistical or classification results. We extracted a fixed 50-sample segment (centered on the burst midpoint, ±25 samples) from the raw beta signal to ensure consistency across bursts of different durations. We then extracted four nonlinear features, including FD, WE, NEO, and SE, from each burst segment. This procedure allowed for a detailed assessment of beta burst dynamics across different temporal scales and conditions, with results stored separately per subject and duration group for further statistical analysis.

A schematic overview of the beta burst detection and duration-based labeling procedure is provided in Fig. 4. The figure illustrates the beta-band EEG signal, the corresponding amplitude envelope, the applied detection threshold, and the identification of individual beta bursts. Burst duration was defined as the time interval during which the envelope exceeded the threshold and was used to classify bursts into short (<100 ms), medium (100–149 ms), and long (≥150 ms) categories prior to nonlinear feature extraction. In addition, a real representative example of beta burst detection, including envelope-based thresholding and the temporal modulation of beta burst probability relative to movement onset, are provided in Supplementary Figure S3.

**Fig. 4.**
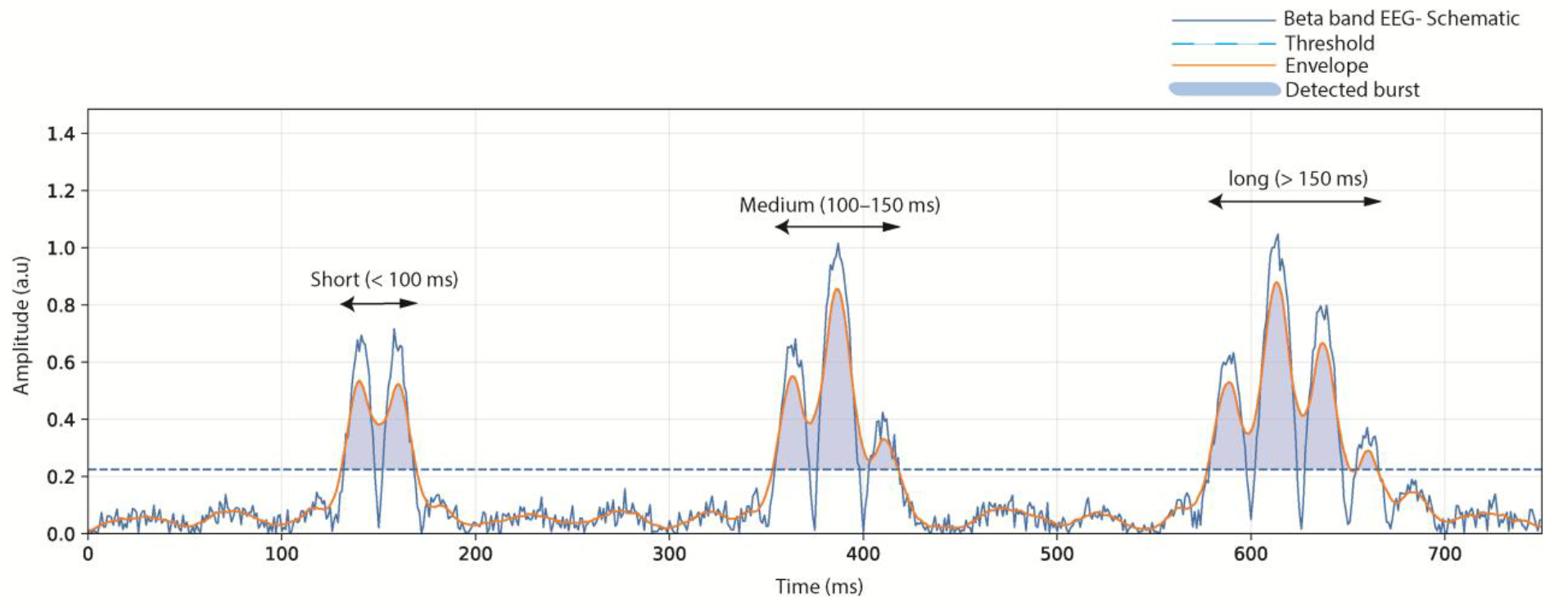
Schematic illustration of beta-burst detection and duration-based labeling. A representative beta-band EEG signal (schematic) is shown together with its amplitude envelope and the applied detection threshold. Shaded regions indicate detected beta bursts, defined as time intervals during which the envelope exceeded the threshold. Burst duration was measured as the continuous time above threshold and used to classify bursts into short (<100 ms), medium (100–149 ms), and long (≥150 ms) categories, as illustrated. The numeric threshold level shown in this schematic is illustrative only; in the actual analysis, beta bursts were identified using a percentile-based threshold (75th percentile of the amplitude distribution) as described in section 2.5.

To assess the robustness of the results to the choice of spatial referencing, a supplementary analysis was performed using Laplacian-referenced EEG. This analysis yielded highly similar beta-burst counts and duration distributions across tasks (Supplementary Table S5).

### 2.6. Nonlinear Dynamic Features

In this study, we extracted FD, WE, NEO, and SE. The rationale for selecting these features and their theoretical underpinnings is detailed in the following subsections.

#### 2.6.1. Fractal Dimension

FD quantifies the complexity and irregularity of a time-series signal across multiple scales. In EEG, particularly within the beta band, FD reflects how dynamically structured neural activity is over time. Higher FD values indicate more intricate and less predictable signal patterns, often associated with increased cortical engagement. During motor tasks, such as isometric contraction or movement initiation, the complexity of beta bursts can vary with motor planning, inhibition, or execution demands. FD was therefore chosen to capture these temporal variations in signal complexity. For instance, during dual-task conditions, cortical processing demands are higher, potentially increasing signal complexity. Thus, FD provides a compact quantitative index for distinguishing sustained from transient beta activity and for assessing how burst duration relates to task-dependent neural dynamics [22]. In the current study, FD was computed using the Higuchi algorithm, with maximum scale parameter was set to 8 (k_Max_ = 8).

#### 2.6.2. Wavelet Entropy

WE quantifies the degree of spectral disorder in a signal by decomposing it into multiple frequency bands using a discrete wavelet transform and assessing how energy is distributed across those bands. WE is particularly suitable for capturing their transient spectral dynamics because beta bursts are short-lived and localized in both time and frequency. A higher WE value indicates a more irregular or broadband energy distribution, whereas lower values suggest more organized, narrowband oscillatory activity. For instance, tasks involving fine motor control (e.g., repetitive finger movements) may yield greater spectral dispersion and higher WE compared to sustained isometric contractions. Accordingly, WE serves as an effective index of how beta-band complexity and spectral organization vary with task context and burst duration [23]. In this study, WE was computed using a Discrete Wavelet Transform (DWT) by decomposing each 50-sample segment into three levels using a Daubechies-4 (db4) mother wavelet. For each level, sub-band energies were calculated from the wavelet coefficients and normalized to obtain a relative energy distribution, after which Shannon entropy was applied to quantify spectral complexity. A 3-level decomposition was selected to balance time–frequency resolution with estimation stability. In very short segments (50 samples), higher decomposition levels substantially reduce the number of coefficients per sub-band approximately proportional to N/2^j^ and increase sensitivity to boundary effects, which can render energy estimates unstable and less physiologically meaningful.

#### 2.6.3. Nonlinear Energy Operator

The instantaneous energy of beta-band EEG activity was quantified using NEO, originally introduced by Kaiser. The NEO captures rapid, local changes in both amplitude and frequency by operating directly on neighboring samples in the time domain.

For a discrete-time signal x(n), the NEO is defined as:

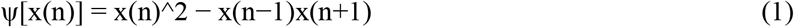

This formulation requires only three consecutive samples and does not rely on window-based statistics, making it particularly well suited for analyzing short, transient oscillatory events such as beta bursts. Because the NEO provides an instantaneous estimate of signal energy, it is largely insensitive to window length and remains stable even when applied to very short EEG segments. In the present study, NEO was computed for each 50-sample burst-centered segment, and the resulting values were averaged within each segment to obtain a single energy-related feature per burst [24].

#### 2.6.4. Sample Entropy

SE measures the irregularity or unpredictability of a time-series signal. Unlike traditional entropy measures such as Approximate Entropy, SE is less sensitive to noise and performs reliably on short data segments, making it well suited to transient EEG analysis. SE is particularly useful for assessing the temporal complexity of EEG signals within and around beta bursts. Changes in burst regularity might reflect task-specific motor control strategies or neural efficiency, for instance, simpler bursts in routine tasks vs. more complex ones in dual-task scenarios. SE helps determine whether beta activity within a task is more predictable (sustained contraction) or irregular (burst-like patterns during movement transitions), making it a strong candidate for burst detection based on duration and temporal structure [25]. In the current study, SE was computed with embedding dimension m=1 and tolerance r=0.3*SD of the signal. These parameters were selected to improve estimation stability on short segments (50 samples), where higher embedding dimensions and lower tolerances can lead to insufficient template matches and unstable entropy values.

Since EEG signals are inherently non-stationary, nonlinear features were computed on short, fixed-length windows to preserve local stationarity and to capture transient dynamics such as beta bursts. FD was originally designed for irregular and non-stationary time series and has been shown to provide stable estimates even for short segments, with performance depending more strongly on algorithmic parameters (e.g. k_Max_) than on window length [26]. SE was introduced to reduce the bias and record-length sensitivity of approximate entropy when applied to short and noisy physiological signals and is commonly used for relative complexity comparisons under fixed windowing [27]. WE was explicitly developed for the analysis of short-duration, non-stationary brain electrical signals and is well suited for transient oscillatory activity [28]. Finally, the NEO is an instantaneous measure of signal energy that operates on only a few consecutive samples and does not rely on long-window statistics [29]. These properties support the use of short analysis windows for consistent nonlinear characterization of beta-burst dynamics.

### 2.7. Statistical Analysis

For each task, the mean values of the four nonlinear features were computed separately for each beta burst duration category (short, medium, and long) on a per-subject basis. This resulted in 26 observations per duration category per task, forming the dataset used for subsequent statistical and ML analyses. To examine how nonlinear features varied with burst duration within each task, one-way ANOVAs were performed across the three duration groups (short, medium, and long) in 3 tasks.

To further investigate the joint influence of burst duration and task condition on each feature, two-way ANOVAs were carried out with task and burst duration as within-subjects factors. Interaction effects were assessed to determine whether burst duration modulated the impact of task conditions on each feature. All statistical tests were two-tailed with a significance threshold of p < 0.05. These analyses aimed to determine whether nonlinear dynamic properties of beta bursts are sensitive to burst duration, task condition, or their interaction.

### 2.8. Feature selection

To gain insights into which features contributed most significantly to classification performance, feature importance was analyzed using the Relief algorithm. Relief is particularly suited for identifying relevant features in noisy and complex datasets like EEG, as it evaluates the ability of each feature to distinguish between neighboring instances of different classes. This analysis helps in understanding the physiological relevance of the selected features (FD, WE, NEO, SE) in relation to beta burst characteristics across tasks [30].

### 2.9. Machine learning

The primary aim of this study was to assess whether nonlinear dynamic features differ across beta-burst duration categories and task conditions at the subject level. So as described earlier, nonlinear feature values were averaged across all detected bursts within each subject and duration category to enable subsequent statistical analysis using repeated-measures ANOVA. Following the statistical validation, an exploratory machine learning (ML) analysis was performed using the same subject-averaged features to assess whether beta-burst duration and task classes could be discriminated based on nonlinear dynamics. The ML analysis was designed as an exploratory step and was not intended to provide a fully optimized or generalized classification framework.

This study conducted a comprehensive comparison of several ML classifiers to empirically identify the most effective models for classifying beta bursts according to their duration and motor task conditions. Because no established consensus exists on the optimal classifier for EEG-based beta burst detection, a broad evaluation was necessary to determine which approaches best capture the discriminative characteristics of nonlinear EEG features. The evaluated classifiers included Decision Tree (DT), Linear Discriminant (LD), Quadratic Discriminant (QD), Logistic Regression (LR), Naïve Bayes (NB), Neural Network (NN), Support Vector Machine (SVM), Random Forests (RF), and k-Nearest Neighbors (k-NN).

Since each observation corresponded to one subject (subject-averaged features), all data splitting and cross-validation were performed at the subject level to prevent subject leakage. First, 15% of subjects were reserved as an independent hold-out test set and were not used during model development. On the remaining 85% of subjects, model development employed a subject-wise grouped 5-fold cross-validation strategy, where folds were constructed such that all data from a given subject were contained entirely within a single fold. In each iteration, four folds were used for training and one fold for validation.

Within each fold, all preprocessing steps (e.g., normalization) and feature selection procedures were fitted exclusively on the training subjects and then applied to the validation subjects. Hyperparameters were optimized within this framework using grid search. For each classifier, mean validation accuracy across the five folds was computed, and the hyperparameter configuration yielding the highest mean accuracy was selected as optimal.

The following hyperparameters were optimized using grid search within the 5-fold CV framework. For the SVM, kernel type (linear or RBF), penalty parameter, and kernel width were explored. The NN models varied in hidden-layer size (5–50 neurons), activation function (tanh or ReLU), and learning rate (0.001–0.01). For RF, the number of trees (50–300) and maximum depth (5–20) were optimized. The DT models were tuned for maximum depth (3–20) and minimum leaf size (1–5). The KNN classifier was optimized for the number of neighbors. The LR models were regularized using the penalty type (L1 or L2) and inverse regularization strength. The LD and QD models required only regularization parameters to stabilize covariance estimation. NB models were trained with Gaussian and kernel variants, and the smoothing parameter was optimized.

After cross-validation, the selected configuration was retrained on the entire 85% training/validation dataset to fully utilize the available data. The resulting model was then evaluated once on the independent 15% test set, which was not used at any point during model development, providing an unbiased estimate of generalization performance. The overall procedure followed Kohavi’s cross-validation framework [31].

Two complementary classification paradigms were implemented: the intra-task paradigm, which differentiated bursts of varying durations within a single motor task, and the inter-task paradigm, which distinguished bursts of the same duration across different motor tasks. Independent models were trained for each scenario to ensure fair comparison across binary and multiclass conditions.

Input features comprised four nonlinear dynamic features (FD, WE, NEO, and SE) with NEO excluded in specific cases based on Relief-based feature relevance analysis. The target variable represented either burst duration (short, medium, or long) or motor task condition. This systematic design enabled an unbiased evaluation of classifier performance and provided insight into which ML models and nonlinear features most effectively differentiate beta bursts across varying temporal and task contexts. The complete processing workflow is illustrated in Fig. 5.

**Fig. 5.**
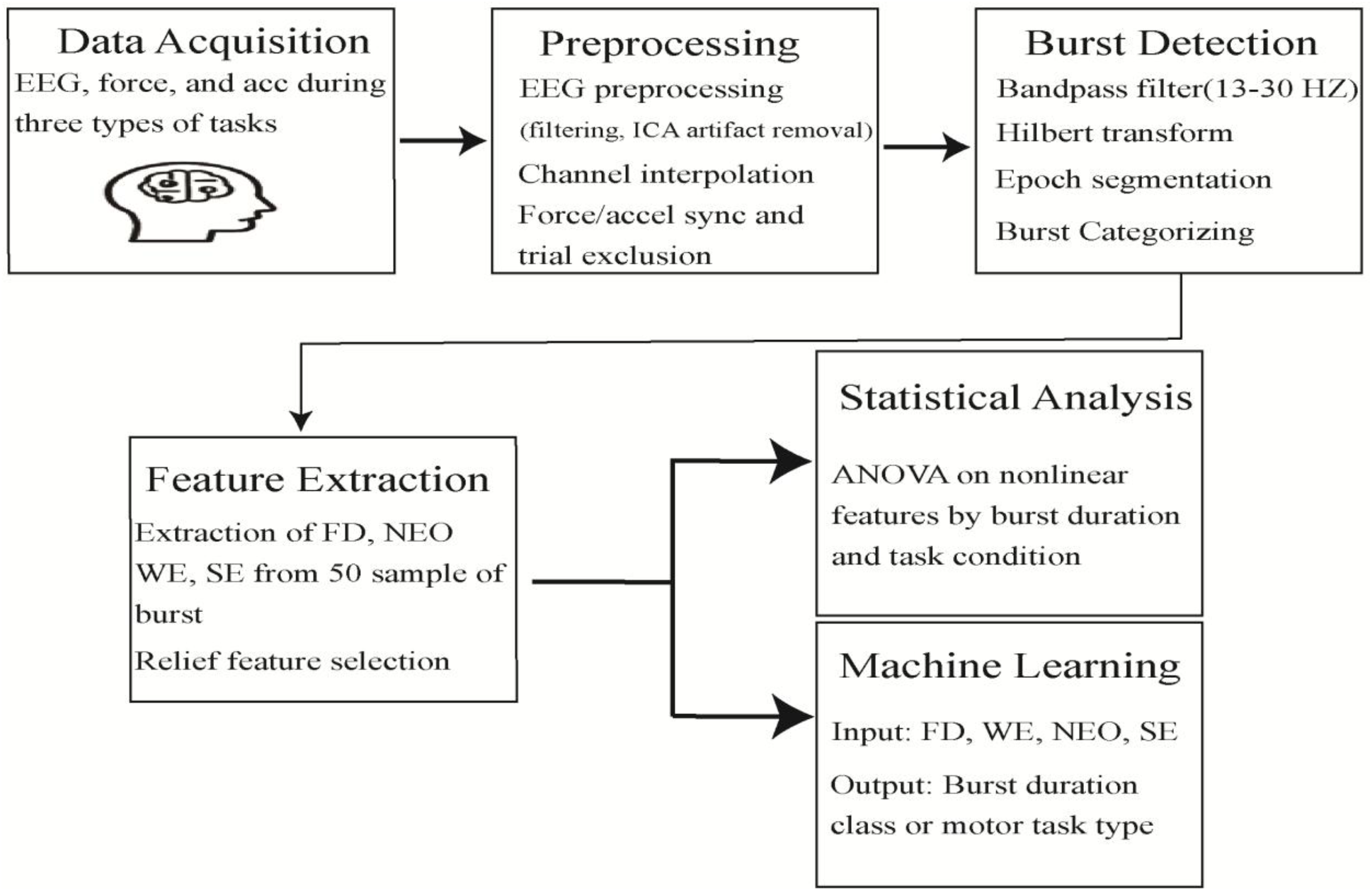
Overview of the EEG beta burst analysis workflow. The diagram summarizes the main stages of the study, including EEG preprocessing, beta burst detection, nonlinear feature extraction, feature selection, and machine learning–based classification.

Model performance was primarily evaluated using classification accuracy because burst duration and task classes were balanced by design at the subject level, and nonlinear features were extracted from fixed-length 50-sample segments centered on each burst. Training, validation, and test sets were constructed to ensure equal representation across burst duration categories and motor task conditions, thereby minimizing class-, task-, and subject-related bias. Under these balanced conditions and given the exploratory scope of the ML analysis, accuracy provides an interpretable and appropriate measure of relative discriminability. To further illustrate class-wise performance, confusion matrices for the best-performing models are provided in the Supplementary Material figure S1 and S2.

All feature extraction and classification procedures were implemented in MATLAB 2024b and executed on a standard desktop computer (Intel i7 CPU, no GPU acceleration). The computation of nonlinear features and subsequent classification required only a few milliseconds per beta burst, confirming that the proposed framework is computationally lightweight and compatible with real-time or online decoding applications.

## 3. Results

### 3.1. Descriptive Statistics of Beta Bursts

More than 1000 beta bursts were detected across all participants and each task condition. These bursts were categorized into three duration classes: short (<100 ms), medium (100–150 ms), and long (>150 ms). The mean and standard deviation of the number of bursts per duration class and task condition are presented in figure 6 (more details in Supplementary Table S1). As shown in the figure, most of the detected bursts were of short duration across all tasks, with the burst count was observed during the isometric contraction task (1055 ± 115.77). In contrast, the number of medium-duration bursts was substantially lower, and long-duration bursts were rare across all tasks. It is worth noting that μ-band activity showed the expected movement-related suppression and qualitatively co-varied with beta activity but was not analyzed further in this study.

**Fig. 6.**
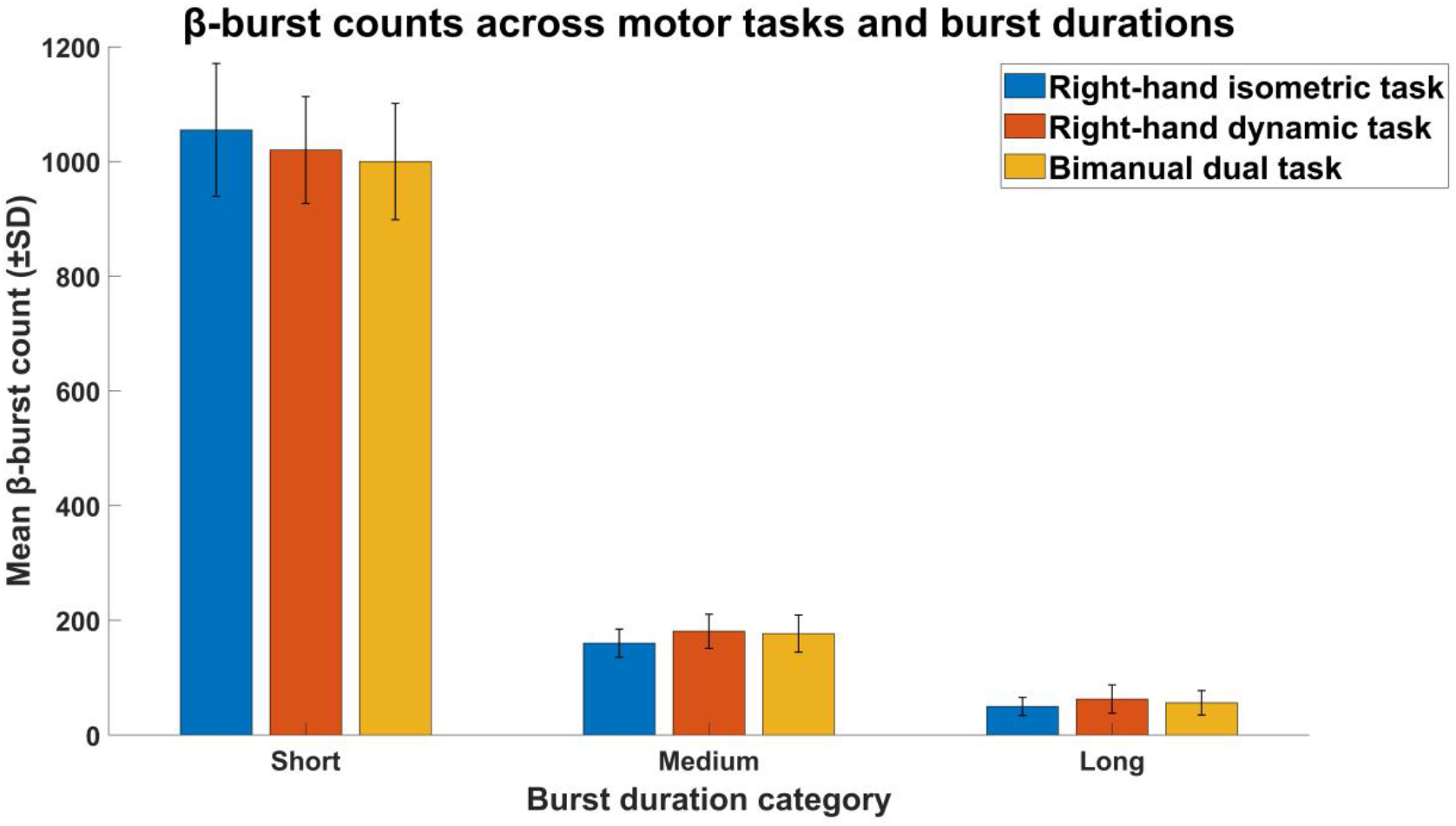
Mean ± SD of β-burst counts across three motor tasks and burst duration categories. Short (< 100 ms) bursts were the most frequent in all conditions, whereas medium (100–150 ms) and long (> 150 ms) bursts occurred progressively less often. Error bars indicate standard deviation across participants.

### 3.2. Group-Level Feature Averages Across Beta Burst Durations and Task Conditions

For each participant, we computed the mean values of four nonlinear features (FD, NEO, SE, and WE) within each beta burst duration category and task condition. These were then averaged across participants to obtain group-level statistics, as detailed in Table 1. This tabular presentation enables a comprehensive comparison of the feature dynamics across tasks and burst durations, offering insights into the variability and consistency of these features across conditions.

**Table 1.** Group-Level mean and standard deviation of FD, NEO, SE, and WE across beta burst categories (short, medium, and long) for three task conditions (isometric contraction, right-hand movement, and dual Task).

|  | Right-hand isometric task |  |  | Right-hand dynamic task |  |  | Bimanual dual task |  |  |
| --- | --- | --- | --- | --- | --- | --- | --- | --- | --- |
|  | Short | Medium | Long | Short | Medium | Long | Short | Medium | Long |
| FD | 1.025±0.0052 | 1.030±0.03 | 1.030±0.006 | 1.025±0.006 | 1.031±0.005 | 1.031±0.007 | 1.026±0.005 | 1.032±0.0042 | 1.031±0.005 |
| NEO | 0.00179±0.005 | 0.0026±0.007 | 0.0026±0.006 | 0.0020±0.0001 | 0.0020±0.0003 | 0.0029±0.0003 | 0.004±0.0005 | 0.005±0.0006 | 0.005±0.0007 |
| SE | 0.27±0.007 | 0.271±0.015 | 0.27±0.019 | 0.276±0.009 | 0.270±0.019 | 0.275±0.025 | 0.27±0.005 | 0.27±0.015 | 0.27±0.020 |
| WE | 0.649±0.0309 | 0.669±0.03 | 0.66±0.05 | 0.658±0.658 | 0.676±0.043 | 0.686±0.077 | 0.656±0.033 | 0.680±0.035 | 0.675±0.40 |

### 3.3. Distributions of Nonlinear Features Across Burst Durations

To evaluate how nonlinear beta burst features varied with burst duration, a one-way ANOVA was conducted for each feature across short, medium, and long categories. The analysis revealed statistically significant effects for several features, indicating that specific nonlinear measures capture duration-dependent burst characteristics.

FD: A significant effect of burst duration was observed, *F*(2, 75) = 9.93, *p* < 0.001, suggesting that FD is highly sensitive to changes in burst length.

SE: SE also varied significantly with duration, *F*(2, 75) = 4.40, *p* = 0.016, reflecting differences in signal irregularity across bursts of different lengths.

WE: WE exhibited a significant main effect, *F*(2, 75) = 3.92, *p* = 0.024, suggesting that frequency–time distribution complexity differs across burst durations.

NEO: No significant effect was found, *F*(2, 75) = 0.13, *p* = 0.881, indicating that instantaneous energy remains relatively stable across burst durations.

These results demonstrate that FD, SE, and WE exhibit duration-dependent modulation, whereas NEO showed no significant modulation with duration, highlighting potential differences in how these nonlinear measures encode temporal burst dynamics.

### 3.4. Distributions of Nonlinear Features Across Task Conditions

To examine how task condition and burst duration jointly influence nonlinear feature distributions, two-way repeated-measures ANOVA were conducted with task condition and burst duration as within-subject factors. The results revealed distinct patterns of sensitivity across features.

FD: A significant main effect of burst duration was observed, *F*(2, 233) = 24.57, *p* < 0.0001, indicating that FD increased with burst duration. Neither the main effect of task condition (*F*(2, 233) = 0.76, *p* = 0.470) nor the interaction between task and duration (*F*(4, 233) = 0.13, *p* = 0.971) was significant, demonstrating that FD is primarily modulated by burst duration, consistent with the earlier one-way ANOVA results.

SE: SE increased modestly with burst duration, *F*(2, 233) = 5.07, *p* = 0.007. A near-significant main effect of task condition was also observed, *F*(2, 233) = 2.32, *p* = 0.050, reflecting slightly higher SE values during right hand dynamic and bimanual dual tasks compared to isometric contraction. No interaction effect was detected (*F*(4, 233) = 0.76, *p* = 0.555), suggesting independent contributions of duration and task.

WE: WE exhibited a significant main effect of burst duration, *F*(2, 233) = 6.00, *p* = 0.003, and a smaller but significant main effect of task condition, *F*(2, 233) = 4.24, *p* = 0.040. No interaction was present (*F*(4, 233) = 0.89, *p* = 0.470). These findings indicate that WE is influenced by both duration and motor task, although the two effects appear additive rather than interactive.

NEO: A significant main effect of task condition was observed, *F*(2, 233) = 6.56, *p* = 0.002, with higher NEO values during dynamic and bimanual tasks compared to the isometric condition. Neither the main effect of burst duration (*F*(2, 233) = 0.20, *p* = 0.822) nor the interaction effect (*F*(4, 233) = 0.10, *p* = 0.983) was significant, suggesting that NEO reflects task-driven rather than duration-dependent changes in signal energy.

Overall, the two-way ANOVA results demonstrate that nonlinear EEG features exhibit distinct sensitivity patterns: FD is predominantly modulated by burst duration, NEO is driven primarily by task condition, and SE and WE show moderate yet independent modulation by both duration and task.

### 3.5. Relief Feature Selection

The Relief algorithm was employed to evaluate the discriminative contribution of each nonlinear feature to the classification of beta bursts [32]. The feature contribution for all classification scenarios are summarized in Supplementary Table S2. Initially, the analysis focused on distinguishing beta bursts by duration within each motor task, considering both binary (two-category) and multiclass (three-category) duration groupings. Subsequently, pairwise comparisons were conducted between bursts of equal duration across different task conditions. As illustrated in figure 7, all four features generally contributed to classification performance. However, in seven specific classification cases, NEO exhibited low relevance scores. In those instances, only the remaining three features (FD, WE, and SE) were retained for model training to optimize classification accuracy.

**Fig. 7.**
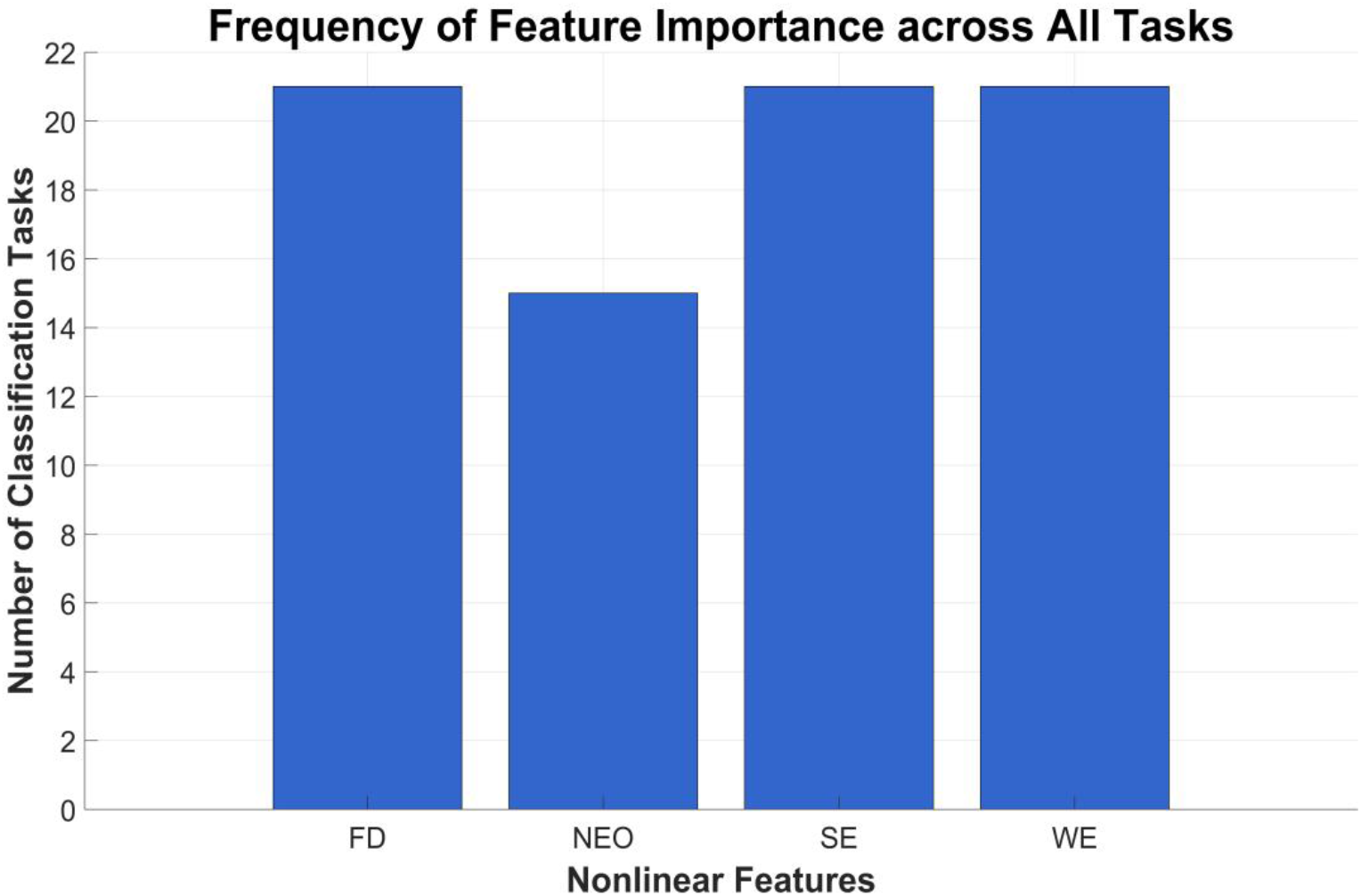
Feature relevance derived from the Relief algorithm. FD, WE, and SE showed the highest discriminative importance for beta-burst classification, while NEO demonstrated lower relevance.

### 3.6. Classification Accuracy Using Machine Learning

The classification performance of multiple ML models was evaluated using a 5-fold cross-validation procedure on 85% of the dataset, with the remaining 15% reserved as an independent test set. Model performance was assessed based on classification accuracy for both validation and test datasets.

Within-task classification by burst duration (figure 8) showed that the optimal model varied depending on the motor task. In Task 1 (right-hand isometric contraction), the QD and DT models achieved the best results, with a validation accuracy of 91.1% and a test accuracy of 85.7% when distinguishing between short–medium and short–long bursts. In Task 2 (right-hand dynamic task), the QD model provided the highest performance for short vs. medium and medium vs. long bursts, while a NN yielded the best test accuracy (85.7%) for short vs. long comparisons. In task 2, when all three duration classes were combined, the QD model achieved a validation accuracy of 71.6% and a test accuracy of 63.6%.

**Fig. 8.**
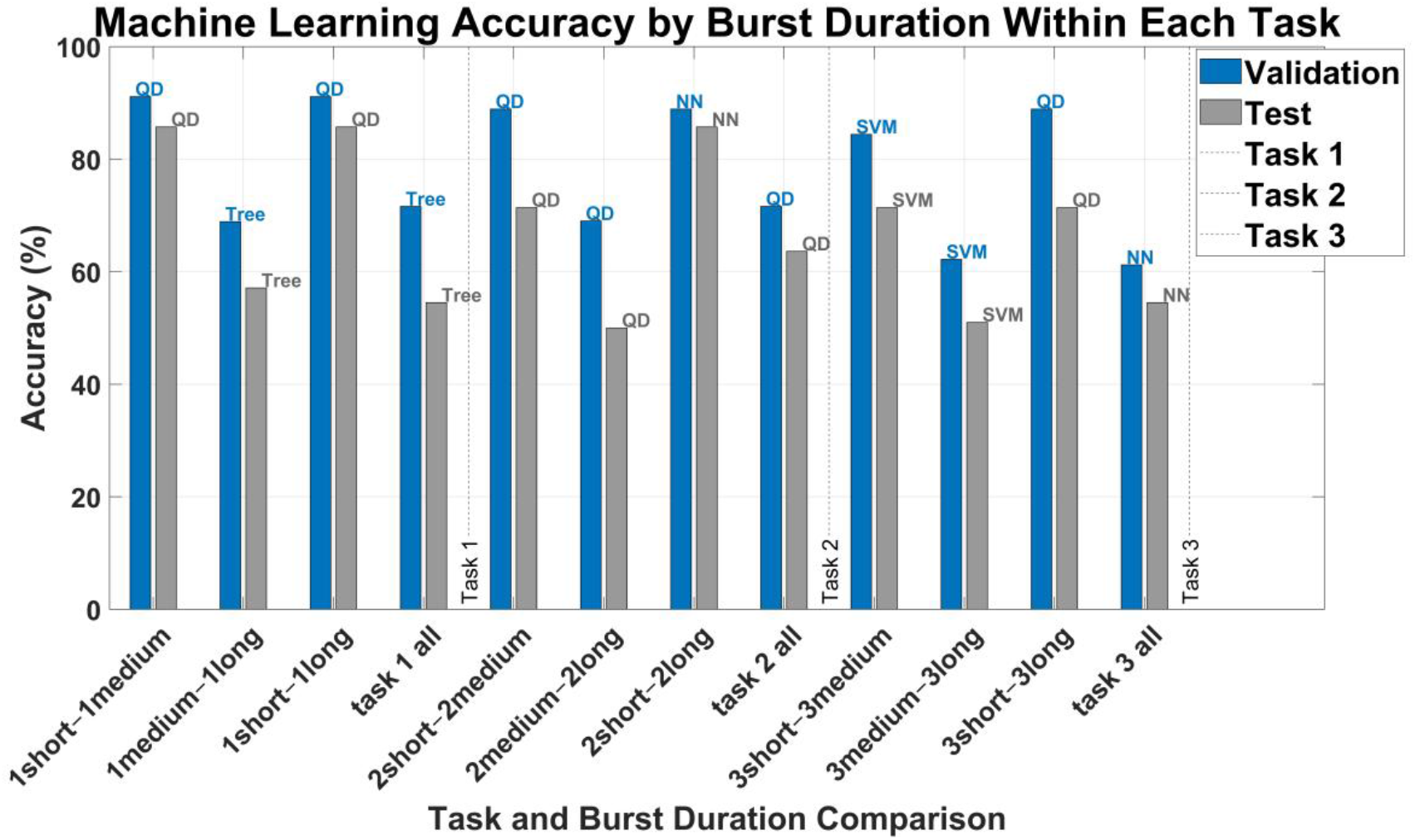
Classification accuracy for beta-burst duration within each motor task. Validation and test accuracies of the best-performing ML models are shown for all duration comparisons (short, medium, long) within the right-hand isometric, right hand dynamic, and bimanual dual tasks. Each bar represents the accuracy performance of the optimal model for that comparison, with associated best models annotated (QD = Quadratic Discriminant, NN = Neural Network, SVM = Support Vector Machine, DT = Decision Tree).

In Task 3 (bimanual dual-task condition), the SVM achieved the highest validation accuracy (84.4%) for short vs. medium bursts, whereas the QD model performed best for short vs. long comparisons. When classifying all three duration categories simultaneously, the NN model achieved moderate results (61.2% validation, 54.5% test), indicating increased difficulty in distinguishing all burst types within a single model (for more detail you can see Supplementary Table S3a).

Cross-task classification (Fig. 9) of bursts with equal duration further highlighted model-specific trends. For short-duration bursts, the DT classifier achieved the best performance when distinguishing between Tasks 2 and 3 (68.9% validation, 57.1% test). For medium-duration bursts, the KNN model achieved high validation (84.4%) and test (71.4%) accuracies. Among long-duration bursts, the SVM model performed best (68.9% validation, 55.2% test) (for more detail you can see Supplementary Table S3b).

**Fig. 9.**
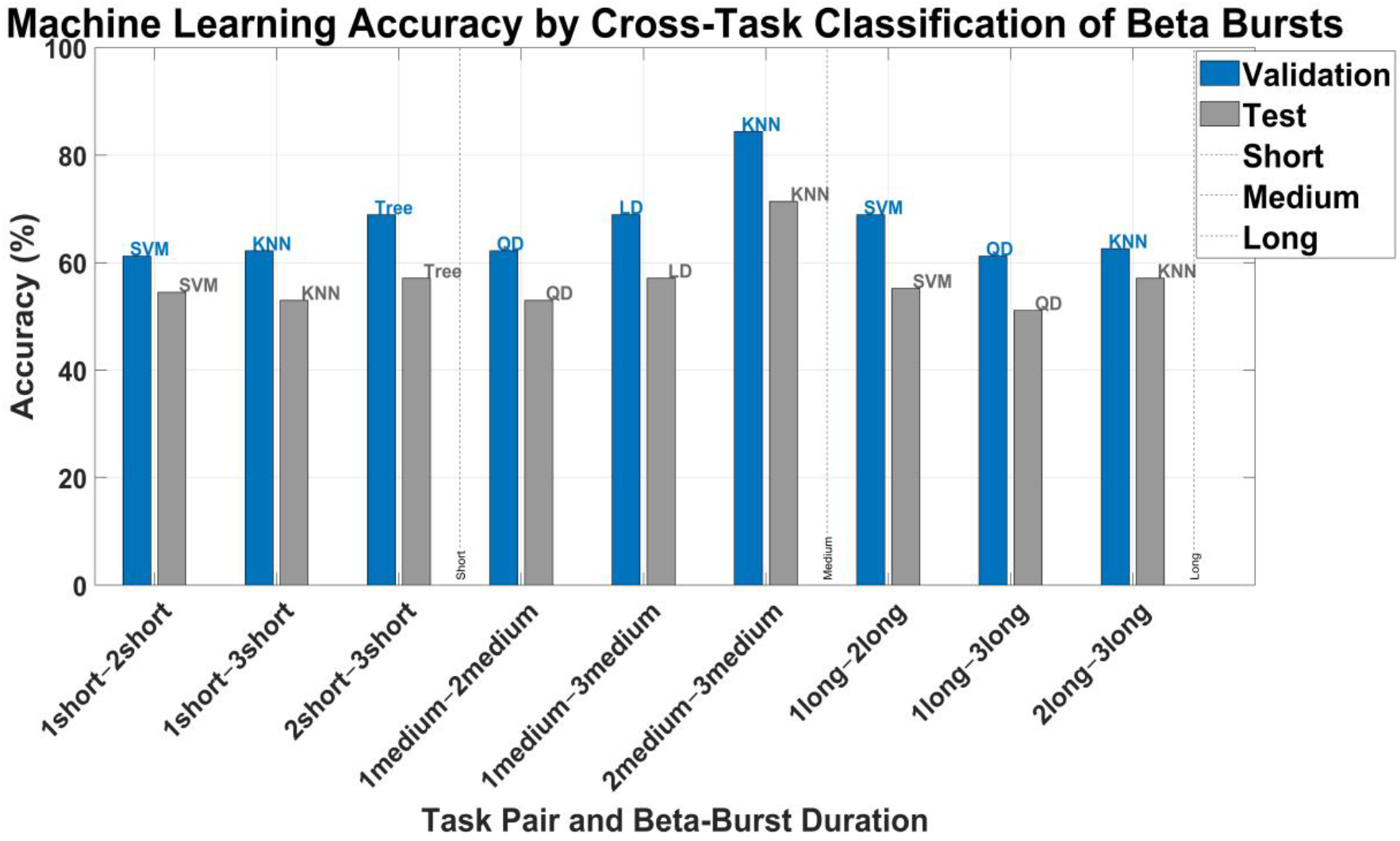
Cross-task classification accuracy of beta bursts with equal duration. Validation and test accuracies of the best-performing ML models for classifying short, medium, and long beta burst across task pairs. Bars indicate accuracy performance for each duration and task comparison, with associated best models annotated (KNN = k-Nearest Neighbor, QD = Quadratic Discriminant, SVM = Support Vector Machine, LD = Linear Discriminant, DT = Decision Tree). Accuracy was highest for medium-duration bursts, indicating stronger generalization across tasks for this category.

Overall, classification accuracy varied across tasks and burst-duration comparisons. The highest accuracies were observed when differentiating between short and medium, as well as short and long, bursts within individual tasks (up to 91.1%), and in pairwise task comparisons for medium bursts (84.4%). In general, ML classifiers exhibited higher accuracy for within-task duration classification than for cross-task comparisons. Optimal performance was achieved by DT, SVM, QD, KNN, and NN models, depending on the specific task and burst-duration configuration.

## 4. Discussion

This study examined whether nonlinear dynamic features including FD, WE, NEO, and SE can effectively characterize and classify beta bursts according to burst duration and motor task condition. The results support the proposed hypotheses: these nonlinear features differed significantly across both burst durations and task types, indicating that they capture structured, task-relevant dynamics in beta activity. Although direct comparisons with conventional linear features (e.g., amplitude or power) were not included, the discriminative performance of nonlinear features alone suggests that they encode complementary or additional information beyond traditional signal metrics.

### 4.1. Methodological Strengths and Innovations

This study introduces several methodological advances that enhance both the analytical rigor and translational potential of the proposed framework.

First, the use of short, fixed-length EEG segments (50 samples per burst) for feature extraction ensures consistent comparisons across bursts of varying durations. This windowed design minimizes the influence of burst length on feature estimation while preserving essential temporal dynamics. Unlike variable-length or template-based burst definitions, fixed-length segmentation improves interpretability and facilitates real-time implementation, where timing precision is critical.

Second, the proposed framework departs from recent high-complexity modelling approaches by offering a lightweight, interpretable, and computationally efficient alternative. For example, Szul et al. [10] employed a biophysical modelling approach to explore beta burst waveform variability, using an adaptive burst detection algorithm and PCA on waveform motifs. While their method provided rich insight into burst diversity, it involved computationally expensive iterative TF decomposition, aperiodic component removal, and PCA modelling, making it difficult to scale for real-time or embedded neurotechnology systems. Likewise, Pauls et al. [11] used wavelet-based amplitude envelope thresholding combined with spectral parameterization (via FOOOF) to decompose beta activity into periodic and aperiodic components in MEG resting-state data. While effective for biomarker identification, that method relies on individualized peak frequency estimation, multi-step artifact correction, and manual channel selection, making it less adaptable to task-related EEG decoding. In contrast, the present study uses a fixed, general-purpose pipeline: we extract four well-known nonlinear dynamic features, FD, WE, NEO, and SE, directly from the time-domain EEG beta band, without requiring spectral fitting, component decomposition, or waveform alignment. These features are task-agnostic, interpretable, and robust, providing meaningful physiological information with minimal computational burden. This approach allows for fast classification using traditional ML models, showing high discriminative power even with small training datasets.

Third, the synchronization of EEG with accelerometer signals in task 2 and 3 ensures that detected beta bursts are tightly aligned with motor behaviour, reducing ambiguity about their functional relevance. This dual-modality integration enhances ecological validity and provides a firm foundation for applying burst-based decoding in motor neuroprosthetics, adaptive deep brain stimulation (DBS), and BCI systems.

Finally, the overall pipeline including, short burst segmentation, simple feature extraction, and supervised classification forms a translational bridge between current neuroscience models and real-time neural interfaces. By avoiding black-box dimensionality reduction or simulation-heavy modelling, this methodology prioritizes speed, reliability, and clinical scalability.

### 4.2. Interpretation of Nonlinear Features in the Context of Motor Control

#### 4.2.1. Fractal Dimension Reflects Cortical Complexity

The significant increase in FD with beta burst duration reflects a key characteristic of fractal-based neural dynamics: longer bursts are not merely extended in time, but structurally more complex. This finding is consistent with earlier studies suggesting that FD captures the self-similarity and scale-free dynamics of EEG, particularly under cognitively or behaviorally engaged states [22, 33]. The present results suggest that FD reflects the temporal richness of neural activity rather than task-specific activation. This supports theoretical models positing that longer beta bursts may reflect prolonged engagement of cortical assemblies or recurrent feedback loops [34], potentially stabilizing motor representations or maintaining postural control. The lack of task modulation implies that FD may serve as a general-purpose marker of intrinsic cortical engagement, regardless of the motor context. These findings align with prior work showing that higher FD is associated with increased information processing demands and complex sensorimotor integration [26]. Collectively, the results imply that FD indexes the dimensional expansion of cortical dynamics as the system transitions from transient to sustained activity.

#### 4.4.2. Sample Entropy Indicates Task-Specific Neural Variability

SE quantifies the degree of unpredictability or irregularity in a time-series signal and is widely applied to assess the temporal complexity of brain activity. In this study, SE showed statistically significant sensitivity to both burst duration and task condition, suggesting that the temporal structure of beta bursts is flexibly modulated by both intrinsic burst characteristics and external motor demands. This dual sensitivity supports theoretical models in which neural variability is not random noise but rather reflects context-dependent adaptation of cortical dynamics. Prior studies have shown that entropy metrics vary with levels of cognitive effort, arousal, and motor control demands [27, 35]. In line with this, our findings suggest that SE may act as a dynamic indicator of the brain’s flexibility or efficiency, adapting in response to both the duration of neural engagement (as indexed by burst length) and the nature of the task being performed.

Notably, the lack of a significant interaction effect between task and duration implies that these two factors exert independent influences on SE. That is, both task context and burst duration contribute to shaping neural regularity but do so through distinct and additive mechanisms. This highlights SE’s potential utility as a feature that can capture both the intrinsic dynamics of the neural signal and the functional context in which it occurs, without being overly dependent on one or the other. Taken together, these findings reinforce the role of SE as a sensitive and interpretable measure of neural complexity, capable of detecting both temporal and functional modulations in beta burst activity across different motor states.

#### 4.2.3. Wavelet Entropy Captures Frequency Complexity Across Tasks

In this study, WE was significantly modulated by both beta burst duration and motor task condition, suggesting that the spectral composition of beta activity is shaped by both the intrinsic properties of the bursts and the functional demands of the task. This indicates that WE is sensitive to multiple dimensions of neural variability: it reflects how the brain flexibly distributes beta power across time and frequency as a function of burst duration and behavioral context.

The absence of a statistically significant interaction between task and duration suggests that these two factors influence WE through independent mechanisms. Longer bursts may recruit broader or more sustained neural populations, producing greater spectral dispersion, whereas task-specific conditions (e.g. static, dynamic, or bimanual movements) likely modulate cortical engagement patterns. Consequently, WE appears to encode both internal burst dynamics and task-driven modulation without one systematically altering the influence of the other.

This interpretation is supported by prior work showing that neural events like beta bursts are not spectrally uniform but instead span a range of frequencies depending on behavioral context. Rosso et al. [28] introduced WE to characterize such short, non-stationary EEG events, finding that greater spectral dispersion often reflects higher cognitive or sensorimotor engagement. Similarly, Cole and Voytek [36] argued that beta oscillations vary markedly in waveform shape and frequency from cycle to cycle, providing a conceptual basis for analyzing beta activity at the burst level rather than assuming uniform spectral structure. Our use of burst-wise entropy directly aligns with this framework, allowing us to capture meaningful frequency variability that traditional band power measures may overlook.

Moreover, the sensitivity of WE to task context aligns with studies using entropy-based features to differentiate between mental or motor states. For example, Subasi [37] showed that spectral entropy measures, including WE, effectively classified EEG signals across different motor-imagery conditions, underscoring their utility for decoding transient brain-state changes.

Taking together, the current and prior evidence converge to identify WE as a robust feature of frequency complexity in beta bursts. One that reflects both neural timescale and behavioral context without confounding the two.

#### 4.2.4. Nonlinear Energy Operator: Limited Utility and Interpretation

In contrast to FD, WE, and SE, the NEO demonstrated limited discriminative capacity across burst durations and task conditions (figure 5 and Supplementary table S1). This suggests that while NEO is sensitive to localized energy fluctuations, a property that makes it useful for transient spike detection or identifying abrupt signal changes, its application to beta burst classification may be constrained by the relatively smoother and oscillatory nature of beta rhythms compared to spike-like events.

This observation aligns with previous studies showing that NEO is most effective for analyzing high-frequency transients such as epileptiform discharges or speech-energy fluctuations [24, 38]. In the case of beta bursts, which are oscillatory and often sustained over tens to hundreds of milliseconds, the energy profile captured by NEO may lack the variability necessary for reliable classification across cognitive or motor contexts. Furthermore, NEO’s minimal contribution within the Relief-based feature selection analysis supports its relatively low discriminative power compared to FD, WE, and SE. Nevertheless, the fact that NEO showed some sensitivity in distinguishing some comparison in particular long bursts across complex motor tasks suggests it may still carry niche utility when combined with other features, particularly in settings requiring high temporal resolution or when capturing onset-related dynamics.

### 4.3. Machine Learning as a Tool for Neural Characterization

The application of ML in this study extends beyond classification performance and serves as a tool for uncovering structured, functionally relevant patterns in transient beta bursts that were identified a priori using a threshold-based detection procedure. The ability of ML algorithms to distinguish beta bursts by duration and task context using nonlinear features supports the view that beta bursts act as context-sensitive markers of cortical processing rather than uniform oscillatory events [6, 39].

An important observation is that classification accuracy was higher for within-task comparisons across burst durations than for cross-task comparisons of equal-duration bursts. This suggests that burst features such as complexity and irregularity vary more systematically with duration than with task type. In other words, how long a burst lasts seems to have a stronger effect on its neural characteristics than what kind of motor task is being performed. This may be because bursts of the same duration can occur across tasks and support different roles depending on the situation, for example, helping to start a movement or maintain it. These findings suggest that while beta bursts are shaped by the tasks they occur in, their core properties are more strongly determined by how long they last, highlighting their role as consistent yet adaptable components of brain activity [40].

From a methodological perspective, the present framework combines simplicity, interpretability, and computational efficiency, offering advantages over more complex approaches. By relying on a small set of well-understood nonlinear features and lightweight classifiers, the framework supports transparent and efficient decoding suitable for real-time and clinical applications where interpretability is essential. By allowing explicit identification of which features drive classification outcomes, the framework supports explainable neural decoding suitable for adaptive deep brain stimulation (aDBS), EEG-based BCIs, and neurorehabilitation systems.

Within this framework, the ML analysis should be viewed as an initial step toward assessing the discriminability of duration and task-dependent beta-burst dynamics rather than as a fully optimized predictive model. By first establishing, through subject-level statistical analysis, that nonlinear features vary systematically with burst duration and task context, the subsequent ML results serve to reinforce the presence of structured and informative patterns in transient beta activity. Given the subject-averaged feature representation and limited dataset size, classification performance is interpreted cautiously and not as evidence of robust cross-subject generalization. Future work with larger datasets will enable event-level modeling and subject-wise validation, allowing more comprehensive evaluation of generalization and the application of more advanced learning architectures.

### 4.4. Theoretical Integration and Broader Neuroscientific Context

The present findings contribute to a growing body of evidence indicating that beta bursts are not uniform phenomena, but vary systematically in duration, morphology, and functional significance [41]. The results strengthen the view that burst duration encodes meaningful information about the underlying cognitive and motor state, consistent with frameworks that conceptualize beta oscillations as modulatory control signals rather than passive background rhythms. By demonstrating that nonlinear dynamic features can reliably differentiate bursts across durations and task contexts, the study provides empirical support for dynamic-systems models of brain function, which emphasize transient, nonstationary, and context-sensitive neural states. This aligns with perspectives that position beta bursts as temporal markers for neural gating, state transitions, and action monitoring in the motor system.

### 4.5. Limitation and Future Directions

Several limitations should be acknowledged. First, the sample size, while adequate for within-subject analysis, may limit generalizability across diverse populations or clinical conditions. Future studies should replicate these results in larger cohorts, including individuals with neurological disorders such as PD and stroke, to evaluate clinical applicability. Second, while we used robust statistical and ML techniques, the initial burst detection relied on a percentile-based threshold, which may still be affected by background signal variability. Future research could explore adaptive or real-time detection algorithms that track nonlinear feature dynamics to identify beta bursts more precisely under varying neural conditions. Third, this study focused on four nonlinear features; incorporating additional features (for example permutation entropy, Lempel-Ziv complexity) or deep learning approaches could further improve classification accuracy. Finally, since the ML analysis was performed on subject-averaged features derived to support the primary statistical objectives, the resulting dataset was relatively small and does not allow conclusions regarding event-level generalization. The ML results should therefore be interpreted as proof of concept, and future studies with larger datasets will explore event-level beta-burst classification using subject-wise validation and hierarchical modeling approaches. Larger datasets would mitigate overfitting, improve cross-subject generalizability, and enable the detection of more nuanced patterns in beta-burst dynamics. Future work should prioritize data expansion both within and across subjects to fully use the potential of ML-based neural characterization.

## 5. Conclusion

This study demonstrates that nonlinear dynamic features, when combined with ML, can effectively classify beta bursts by duration and motor task context, revealing their structured and task-dependent nature. The findings show that nonlinear approaches capture meaningful distinctions in neural activity that conventional methods may overlook, supporting a shift toward more dynamic and functionally grounded models of motor control. By integrating advanced signal processing with scalable, interpretable ML techniques, this work offers a novel and robust approach to analyzing transient neural events. These insights have important implications for both neuroscience research and applied neurotechnology, paving the way for more responsive and real-time neural analysis systems.

## Supplementary Materials

**Figure S1.**
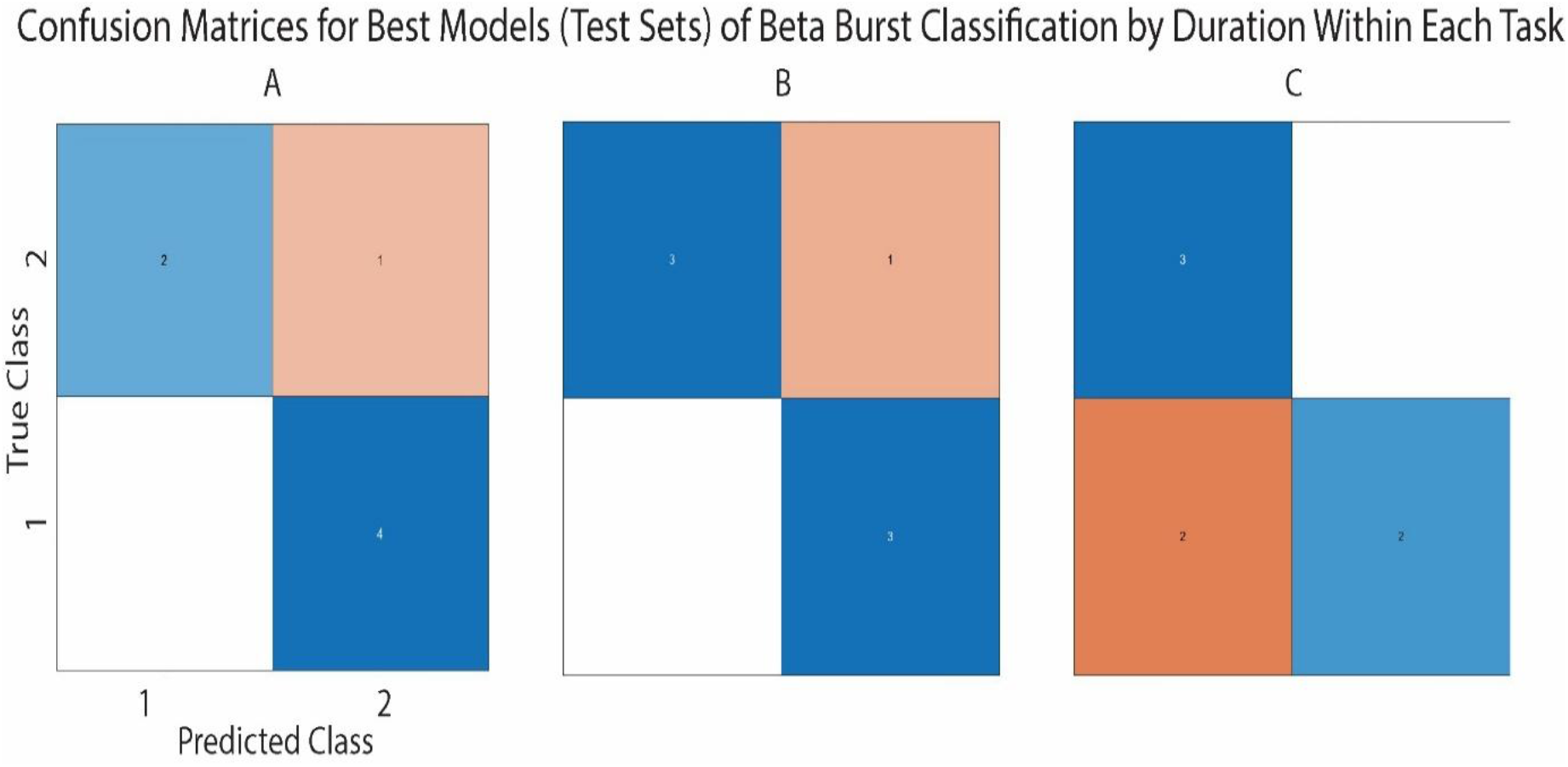
Confusion matrices for intra-task beta-burst classification. Each matrix displays the distribution of true versus predicted burst-duration categories. **A)** Task 1 – right-hand isometric contraction, classified using a QD model (85.7 % test accuracy). **B)** Task 2 – right-hand dynamic movement, classified using a NN model (85.7 % test accuracy). **C)** Task 3 – bimanual dual-task condition, classified using a QD model (71.4 % test accuracy).

**Figure S2.**
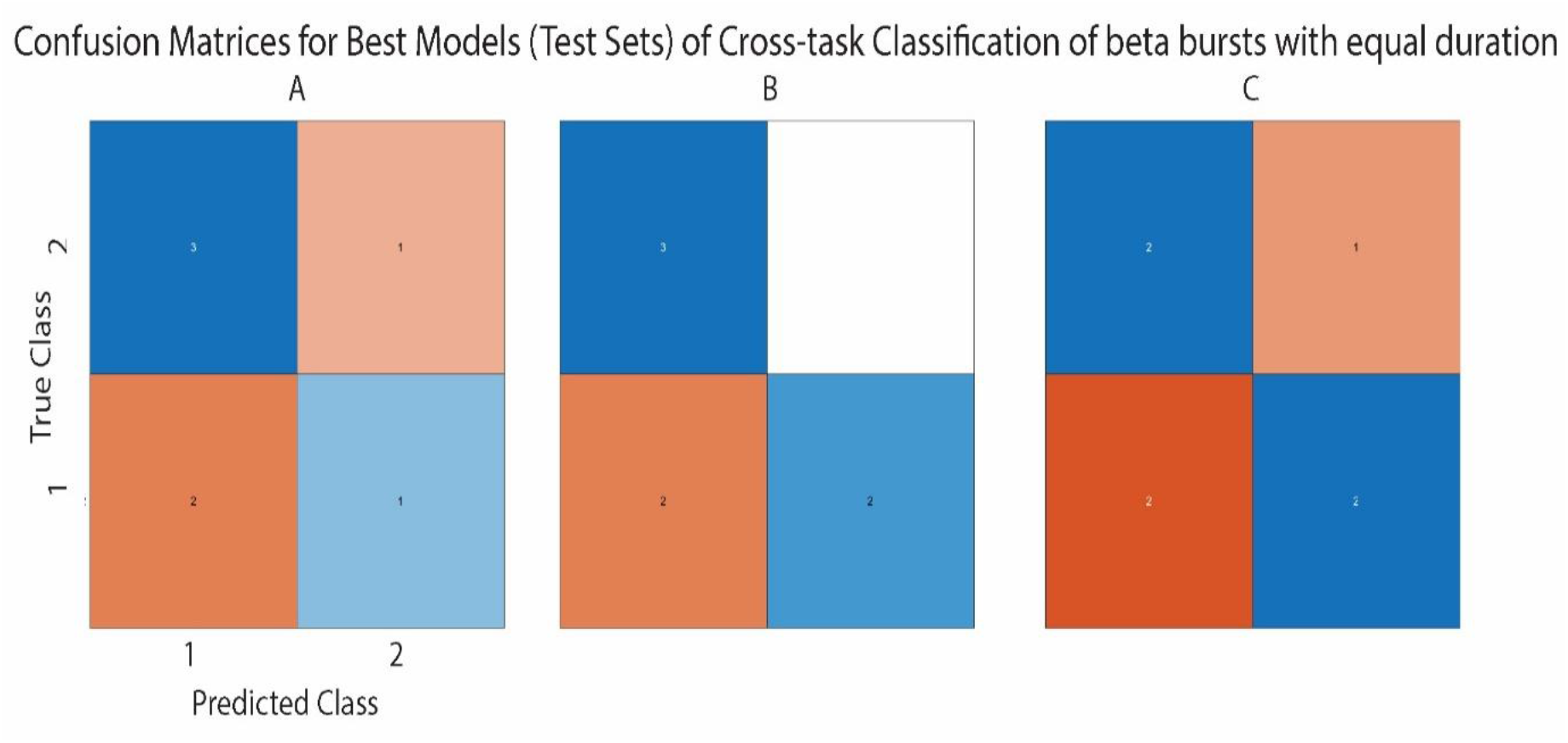
Confusion matrices for cross-task beta-burst classification of equal-duration bursts. Each matrix compares true and predicted task labels, illustrating model generalization across motor-task conditions. **A)** Short-duration bursts, best classified by a DT model (57.1 % test accuracy). **B)** Medium-duration bursts, best classified by a KNN model (71.4 % test accuracy). **C)** Long-duration bursts, best classified by a KNN model (57.1% test accuracy).

**Figure S3.**
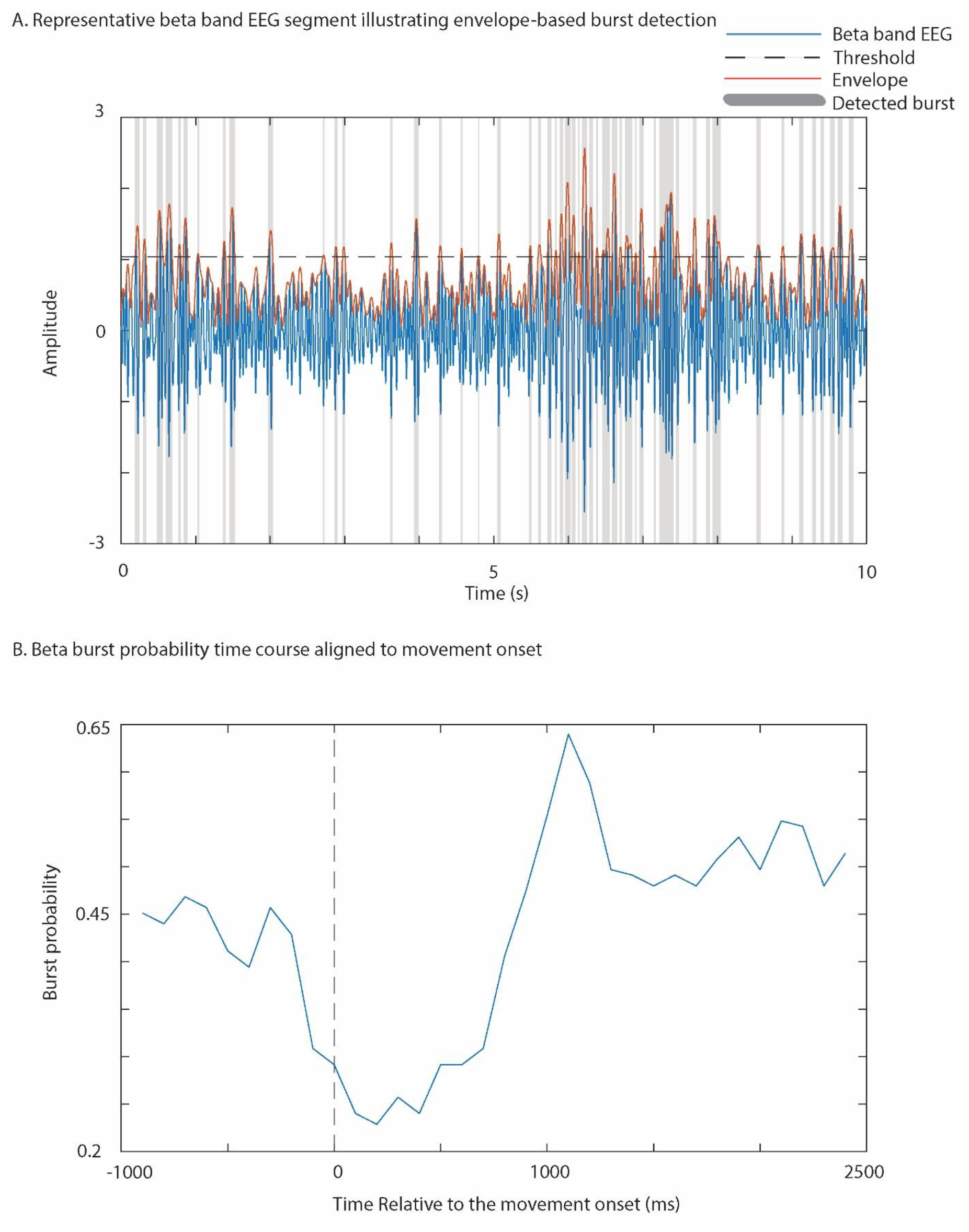
Visualization of beta burst detection and motor-related burst dynamics. (A) Representative 10-s segment of beta band EEG from left sensorimotor electrodes showing the filtered signal, its amplitude envelope, the detection threshold, and detected beta bursts. Bursts were defined as periods during which the envelope exceeded the threshold. Shaded regions indicate detected beta bursts. (B) Time course of beta burst probability aligned to movement onset (0 ms).

**Table S1.** Mean ± SD of beta burst counts across different tasks and burst durations (short duration: <100 ms; medium: 100–150 ms; long: >150 ms)

|  | Right-hand isometric task | Right-hand dynamic task | Bimanual task |
| --- | --- | --- | --- |
| Short | 1055± 115.77 | 1020±93.25 | 1000± 101.26 |
| Medium | 160.30±24.37 | 181±29.69 | 176.92± 32.36 |
| Long | 50±15.75 | 62.73±24.60 | 56.34±21.22 |

**Table S2.** Relief-Based Feature Importance for Beta Burst Classification by Duration and Task.

| Classification Task | FD | NEO | SE | WE |
| --- | --- | --- | --- | --- |
| 1short vs 1medium | ✓ | ✓ | ✓ | ✓ |
| 1medium vs 1long | ✓ | ✗ | ✓ | ✓ |
| 1short vs 1long | ✓ | ✓ | ✓ | ✓ |
| All beta durations in Task 1 | ✓ | ✓ | ✓ | ✓ |
| Taks 2 |  |  |  |  |
| 2short vs 2medium | ✓ | ✓ | ✓ | ✓ |
| 2medium vs 2long | ✓ | ✗ | ✓ | ✓ |
| 2short vs 2long | ✓ | ✓ | ✓ | ✓ |
| All beta durations in Task 2 | ✓ | ✓ | ✓ | ✓ |
| Task 3 |  |  |  |  |
| 3short vs 3medium | ✓ | ✗ | ✓ | ✓ |
| 3medium vs 3long | ✓ | ✗ | ✓ | ✓ |
| 3short vs 3long | ✓ | ✗ | ✓ | ✓ |
| All beta durations in Task 3 | ✓ | ✗ | ✓ | ✓ |
| 1short vs 2short | ✓ | ✗ | ✓ | ✓ |
| 1short vs 3short | ✓ | ✓ | ✓ | ✓ |
| 2short vs 3short | ✓ | ✓ | ✓ | ✓ |
| 1medium vs 2medium | ✓ | ✓ | ✓ | ✓ |
| 1medium vs 3medium | ✓ | ✓ | ✓ | ✓ |
| 2medium vs 3medium | ✓ | ✓ | ✓ | ✓ |
| 1long vs 2long | ✓ | ✓ | ✓ | ✓ |
| 1long vs 3long | ✓ | ✓ | ✓ | ✓ |
| 2long vs 3long | ✓ | ✓ | ✓ | ✓ |

**Table S3a.**
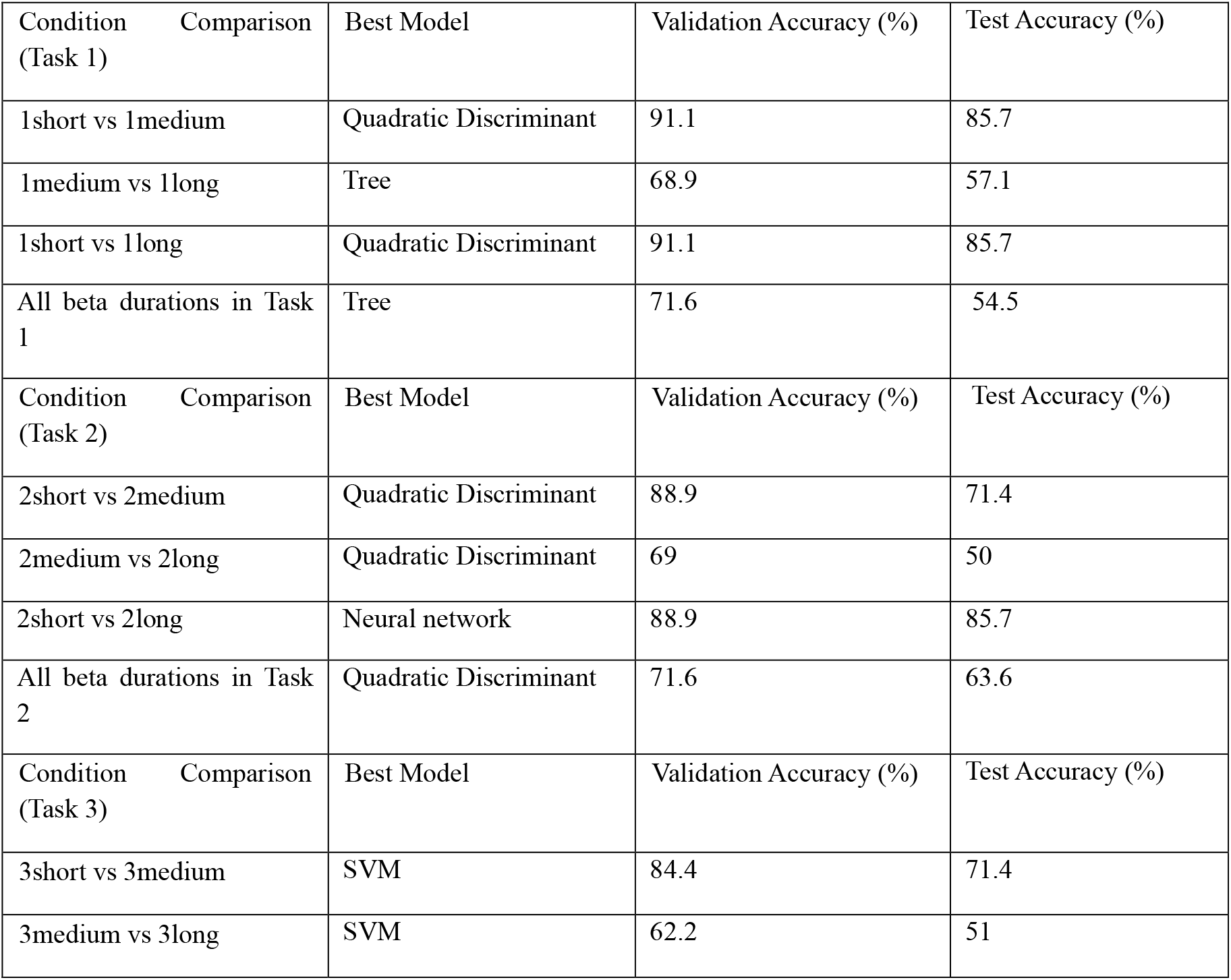
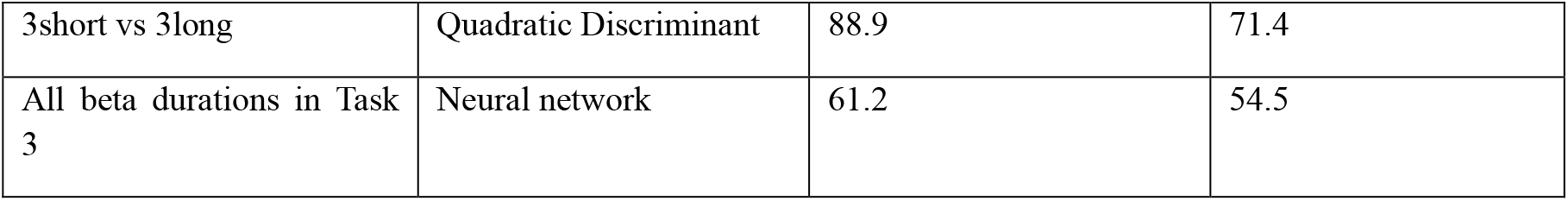
ML Accuracy for Beta Burst Classification by Duration Within Each Task.

**Table 3b.**
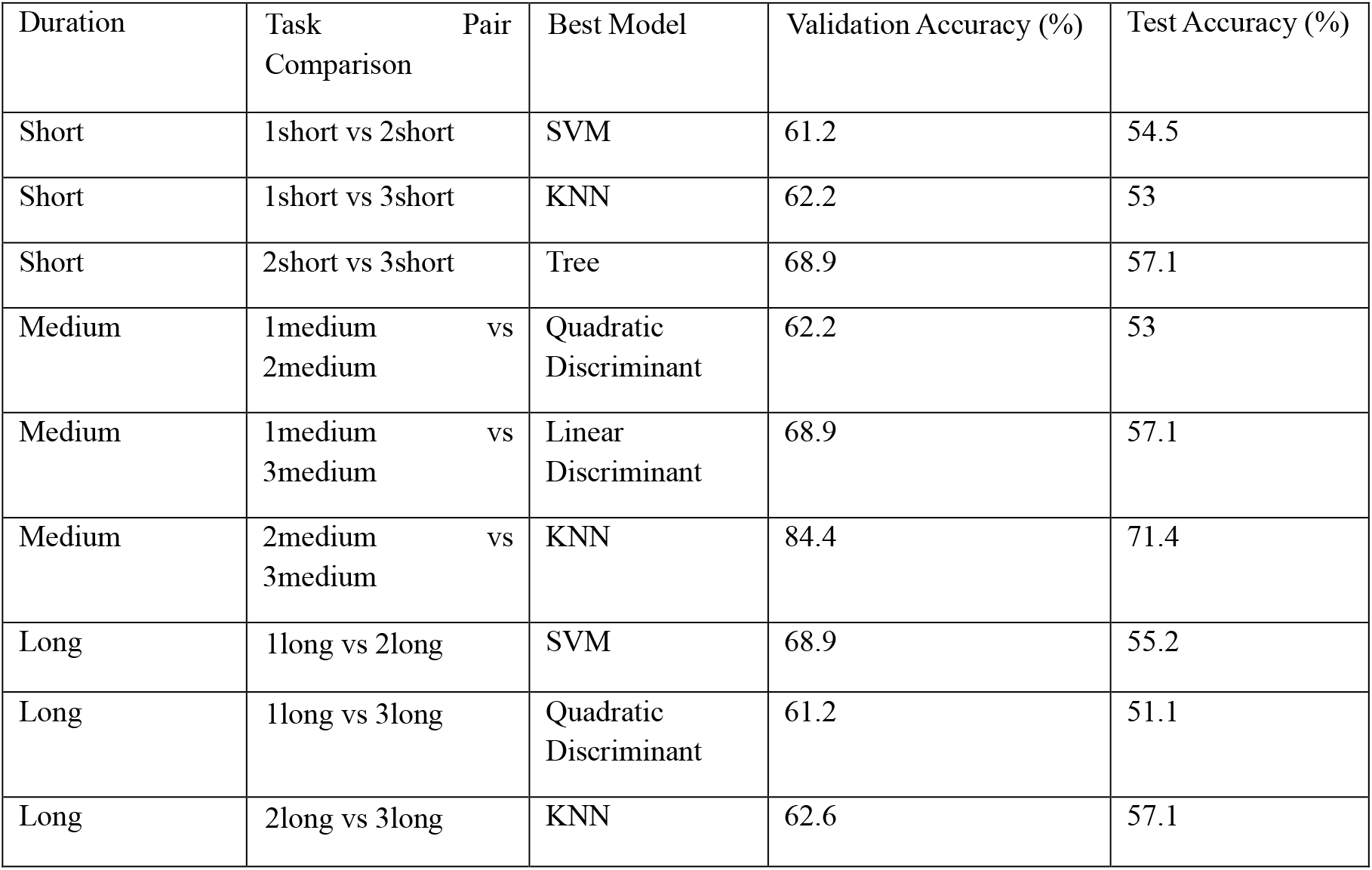
ML Accuracy for Classification of Beta Bursts with Same Duration Across Tasks (Pairwise)

**Table S4.** Mean validation accuracy (%) of all evaluated classifiers averaged across all classification scenarios.

| Model | Mean Validation Accuracy (%) |
| --- | --- |
| DT | 66.69 ± 11.62 |
| RF | 64.51 ± 9.93 |
| QD | 70.57 ± 12.72 |
| SVM | 68.12 ± 10.74 |
| NN | 65.22 ± 12.15 |
| KNN | 65.80 ± 10.97 |
| NB | $60.27 \pm 8.81$ |
| LR | $50.16 \pm 15.70$ |
| LD | $64.69 \pm 10.60$ |

**Table S5.** Mean ± SD of beta-burst counts across different tasks and duration categories using Laplacian-referenced EEG.

|  | Right-hand isometric task | Right-hand dynamic task | Bimanual task |
| --- | --- | --- | --- |
| Short | $955 \pm 85.77$ | $1100 \pm 101.83$ | $962 \pm 120.99$ |
| Medium | $170.70 \pm 20.6$ | $175 \pm 22.03$ | $159 \pm 29.37$ |
| Long | $55 \pm 10.06$ | $61.07 \pm 25.86$ | $52.48 \pm 18.18$ |

